# Herbicidal Quassinoids Isolated from *Ailanthus altissima* Leaves

**DOI:** 10.1101/2023.11.10.566620

**Authors:** Young Sook Kim, Surk-Sik Moon, Jung Sup Choi

## Abstract

Bioassay-guided fractionation of the methanolic extract of *Ailanthus altissima* leaves led to the isolation of one new quassinoid, named 6-α-tigloyloxyailanthone (**3**), and three known quassinoids, ailanthone (**1**), 13, 18-dehydroglaucarubinone (**2**), and 6-α-tigloyloxychaparrinone (**4**) by a series of chromatographic methods. The structures of the isolates were established by one-dimensional (1D) and 2D-nuclear magnetic resonance (NMR) analysis along with high-resolution time-of-flight mass spectrometry (HRTOFMS) and chemical methods. In addition, herbicidal activity against five grassy weeds and five broad-leaf weeds was evaluated.

## Introduction

Quassinoids are a group of degraded triterpenes found in the family Simaroubaceae that undergo extensive oxidative biodegradation, leaving their carbon skeleton highly oxygenated[1-4]. Quassinoids are classified into five groups according to their basic structure, C-18, C-19, C-20, C-22, and C-25. The C-20 quassinoids have been extensively investigated due to their antileukemic activity discovered in the early 1970s[3, 5] Since 1960, hundreds of quassinoids have been isolated and identified from plants, but they seem harder to synthesize due to the presence of their highly oxygenated carbon framework[2]. Many quassinoids display a wide range of biological activities *in vitro* or *in vivo*, including antitumor[6], antimalarial[7], antiviral[8], anti-inflammatory[9], antifeedant[10], insecticidal[11], antiulcer[12], and herbicidal activities[13]. In our ongoing investigations of structurally unique bioactive agents, such as herbicides derived from traditional plants, systematic phytochemical studies of *Ailanthus altissima* resulted in the isolation of one new quassinoid (**3**) and three known quassinoids (**1, 2**, and **4**). In this paper, we describe the isolation, structural elucidation, and evaluation of the herbicidal activities of all isolates obtained.

## Experimental details

### Plant material

Seeds of 10 weed species (*S. bicolor, E. crus-galli, A. smithii, D. sanguiinalis, P. dichotomiflorum, S. nigrum, A. indica, A. avicennae, X. strumarium, C. japonica*) were germinated in flats in a commercial greenhouse substrate and watered with tap water. The weeds were grown in a greenhouse at 30 ± 3 / 20 ± 3 ºC day/night temperature with a 14-h photoperiod.

### Biological activity

The activity assay was performed 12 days after sowing for the foliar application of samples at each concentration with a laboratory spray gun. The herbicidal activity of the foliar application was evaluated by visual injury 14 days after treatment (0, no damage; 100, complete control).

### Extraction and isolation

Fresh samples (5 kg) were dipped in MeOH at room temperature and filtered after three days. After concentrating the methanol, the extract was defatted with hexane, then partitioned between EtOAc and H_2_O. A portion of the active EtOAc fraction (6 g) was purified by gradient elution on a flash silica gel column using CHCl_3_:MeOH (50:1, 20:1, 10:1, 5:1, and 100% MeOH). Fraction II [1.03 g, CHCl_3_-MeOH (20:1)] and fraction III [1.32 g, CHCl_3_:MeOH (10:1)], which were active in the bioassay, were further purified by ODS Sep-pak cartridge (Alltech, Deerfield, IL, USA) chromatography eluted with an increasing methanol concentration gradient (0% – 100%) in water. The active fraction was finally purified by reversed-phase HPLC. Preparative HPLC (C_18_, 5 *μ*m, 20 × 250 mm; COSMOSIL) was performed using 30 – 50% aqueous MeOH, UV detection at 254 nm, a flow rate of 12 mL/min, and gradient elution for 50 min.

### Structural analysis

Melting points were measured on a Fisher melting point apparatus and uncorrected. Optical rotations were measured on a Perkin Elmer 341-LC polarimeter. NMR spectra were recorded on a Bruker 900 and 700 spectrometer with standard pulse sequences, operated at 900 and 700 MHz for ^1^H NMR and 226 and 176 MHz for ^13^C NMR. Chemical shifts, measured in ppm, were referenced to solvent peaks (δ_H_ 2.50 and δ_C_ 39.5 for DMSO-*d*_***6***_). HR-TOF-MS spectra were recorded in the positive ESI mode on a Waters Synapt G2 at the Korea Basic Science Institute.

#### Compound 1 (ailanthone)

^1^H NMR (DMSO-*d*_6_, 900 MHz) δ: 8.23 (1H, s, 11-OH), 7.08 (1H, d, *J* = 2.8 Hz, 1-OH), 5.99 (1H, d, *J* = 0.9 Hz, H-3), 5.40 (1H, d, *J* = 4.5 Hz, 12-OH), 5.05 (1H, d, *J* = 0.9 Hz, H_a_-21), 5.02 (1H, d, *J* = 0.9 Hz, H_b_-21), 4.55 (1H, t, *J* = 2.8 Hz, H-7), 4.28 (1H, s, H-1), 3.81 (1H, d, *J* = 9.0 Hz, H_a_-20), 3.27 (1H, d, *J* = 9.0 Hz, H_b_-20), 3.67 (1H, d, *J* = 3.6 Hz, H-12), 2.88 (1H, d, *J* = 10.8 Hz, H-5), 2.98 (1H, dd, *J* = 18.0, 14.4 Hz, H_a_-15), 2.81 (1H, s, H-9), 2.77 (1H, dd, J = 13.5, 5.4, H-14), 2.44 (1H, dd, *J* = 18.0, 5.4 Hz, H_b_-15), 2.03 (2H, m, H-6), 1.93 (3H, s, H-18), 1.06 (3H, s, H-19) ppm; ^13^C NMR (225 MHz, DMSO-*d*_6_): 197.1 (C2), 169.1 (C16), 162.4 (C4), 146.6 (C13), 125.0 (C3), 117.6 (C21), 108.8 (C11), 82.4 (C1), 79.0 (C12), 77.5 (C7), 71.1 (C20), 46.1 (C14), 44.5 (C8 and C10), 43.3 (C9), 41.2 (C5), 34.3 (C15), 25.1 (C6), 22.4 (C18), and 9.5 (C19) ppm; HRTOFMS (positive ESI mode) *m/z* 399.1415 [M + Na]^+^ (calcd for C_20_H_24_O_7_+Na, 399.1420).

#### Compound 2 (13, 18-dehydroglaucarubinone)

^1^H NMR (DMSO-d6, 700 MHz) δ: 8.33 (1H, s, 11-OH), 7.16 (1H, d, *J* = 2.1 Hz, 1-OH), 5.99 (1H, q, *J* = 0.7 Hz, H-3), 5.64 (1H, br s, H-15), 5.35 (1H, br s, 12-OH), 5.11 (1H, s, 2’-OH), 5.09 (1H, d, *J* = 1.4 Hz, Ha-21), 4.99 (1H, br s, H_b_-21), 4.70 (1H, t, *J* = 2.8 Hz, H-7), 4.42 (1H, d, *J* = 2.1 Hz, H-1), 3.81 (1H, d, *J* = 8.4 Hz, H_a_-20), 3.67 (1H, d, *J* = 4.9 Hz, H-12), 3.32 (1H, dd, *J* = 8.4 Hz, H_b_-20), 3.08 (1H, d, *J* = 11.9 Hz, H-5), 3.04 (1H, d, *J* = 11.4 Hz, H-14), 2.97 (1H, s, H-9), 2.06 (2H, m, H-6), 1.93 (3H, s, H-18), 1.70 (1H, dq, *J* = 140.0, 7.0 Hz, H_a_-3’), 1.53 (1H, dq, *J* = 140.0, 7.0 Hz, H_b_-3’), 1.30 (3H, s, H-5’), 1.06 (3H, s, H-19), 0.81 (3H, t, *J* = 7.0 Hz, H-4’) ppm; ^13^C NMR (176 MHz, DMSO-*d*_6_): 197.0 (C2), 174.2 (C1’), 166.7 (C16), 162.4 (C4), 142.4 (C13), 124.8 (C3), 119.8 (C21), 108.6 (C11), 82.1 (C1), 78.7 (C12), 77.8 (C7), 73.9 (C2’), 70.7 (C20), 68.2 (C15), 50.2 (C14), 46.4 (C8), 44.5 (C10), 44.0 (C9), 40.7 (C5), 32.8 (C3’), 25.8(C5’), 24.6 (C6), 22.2 (C18), 9.5 (C19), and 8.0 (C4’) ppm; HRTOFMS (positive ESI mode) *m/z* 499.1944 [M + Na]^+^ (calcd for C_25_H_32_O_9_+Na, 499.1944) and *m/z* 975.3983 [2M + N]^+^ [calcd for (C_25_H_32_O_9_)_2_+Na, 975.3990].

#### Compound 3 (6-α-tigloyloxyailanthone)

^1^H NMR (DMSO-*d*_6_, 700 MHz) δ: 8.15 (1H, s, 11-OH), 7.28 (1H, d, *J* = 2.8 Hz, 1-OH), 6.91 (dq, *J* = 7.7, 2.1 Hz, H3’), 6.02 (1H, s, H-3), 5.55 (1H, d, *J* = 4.2 Hz, 12-OH), 5.49 (1H, dd, *J* = 11.9, 2.8 Hz, H-6), 5.02 (1H, d, *J* = 1.4 Hz, H_a_-21), 5.01 (1H, s, H_b_-21), 4.63 (1H, d, *J* = 2.8 Hz, H-7), 4.39 (1H, s, H-1), 3.91 (1H, d, *J* = 8.4 Hz, H_a_-20), 3.69 (1H, t, *J* = 4.9 Hz, H-12), 3.46 (1H, d, *J* = 11.9 Hz, H-5), 3.39 (1H, dd, *J* = 8.4 Hz, H_b_-20), 3.01 (1H, dd, *J* = 18.4, 13.3 Hz, H_a_-15), 2.85 (1H, dd, *J* = 13.3, 5.6 Hz, H-14), 2.83 (1H, s, H-9), 2.47 (1H, dd, *J* = 18.4, 5.6 Hz, H_b_-15), 1.94 (3H, s, H-18), 1.829 (3H, s, H5’), 1.823 (3H, d, *J* = 4.2 Hz, H4’), 1.21 (3H, s, H-19) ppm; ^13^C NMR (176 MHz, DMSO-*d*_6_): 196.6 (C2), 168.2 (C16), 165.9 (C1’), 161.2 (C4), 146.1 (C13), 139.2 (C3’), 127.9 (C2’), 127.6 (C3), 117.5 (C21), 109.0 (C11), 82.4 (C1), 79.1 (C12), 77.5 (C7), 70.2 (C20), 67.3 (C6), 47.2 (C10), 46.1 (C14), 45.2 (C8), 44.1 (C5), 42.0 (C9), 34.0 (C15), 24.7 (C18), 14.4 (C4’), 11.9 (C5’), and 10.9 (C19) ppm; HRTOFMS (positive ESI mode) *m/z* 497.1785 [M + Na]^+^ (calcd for C_25_H_30_O_9_+Na, 497.1788) and *m/z* 971.3670 [2M + N]^+^ [calcd for (C_25_H_30_O_9_)_2_+Na, 971.3678].

#### Compound 4 (6-a-tigloyloxychaparrinone)

^1^H NMR (DMSO-d_6_, 700 MHz) δ: 8.0 (1H, s, 11-OH), 7.19 (1H, d, *J* = 2.8 Hz, 1-OH), 6.91 (dq, *J* = 7.7, 2.1 Hz, H3’), 6.0 (1H, s, H-3), 5.49 (1H, dd, *J* = 11.9, 2.8 Hz, H-6), 5.14 (1H, d, *J* = 4.9 Hz, 12-OH), 4.56 (1H, d, *J* = 2.8 Hz, H-7), 4.30 (1H, d, *J* = 2.8 Hz, H-1), 3.93 (1H, d, *J* = 8.4 Hz, H_a_-20), 3.64 (1H, dd, *J* = 8.4 Hz, H_b_-20), 3.39 (1H, d, *J* = 11.9 Hz, H-5), 3.21 (1H, t, *J* = 4.9 Hz, H-12), 2.72 (1H, dd, *J* = 18.2, 13.3 Hz, H_a_-15), 2.61 (1H, s, H-9), 2.40 (1H, dd, *J* = 18.2, 5.6 Hz, H_b_-15), 2.09 (1H, m, H-13), 2.05 (1H, m, H-14), 1.94 (3H, s, H-18), 1.82 (3H, br s, H5’), 1.81 (3H, d, *J* = 7.7 Hz, H4’), 1.22 (3H, s, H-19), 0.85 (3H, d, *J* = 7.0 Hz, H-21) ppm; ^13^C NMR (176 MHz, DMSO-*d*_6_): 196.5 (C2), 168.9 (C16), 165.8 (C1’), 161.2 (C4), 139.0 (C3’), 127.8 (C2’), 127.5 (C3), 109.1 (C11), 82.5 (C1), 78.1 (C12), 77.6 (C7), 69.3 (C20), 67.3 (C-6), 47.1 (C10), 45.7 (C8), 44.1 (C5), 41.8 (C9), 40.6 (C14), 30.2 (C13), 29.3 (C15), 24.5 (C18), 14.3 (C4’), 12.5 (C21), 11.8 (C5’), and 10.8 (C19) ppm; HRTOFMS (positive ESI mode) *m/z* 499.1944 [M + Na]^+^ (calcd for C_25_H_32_O_9_+Na, 499.1944) and *m/z* 975.3983 [2M + N]^+^ [calcd for (C_25_H_32_O_9_)_2_+Na, 975.3990].

## Results and Discussion

The MeOH extract of *Ailanthus altissima* leaves at 10 mg/mL showed 100% control of *Sorghum bicolor, Digitaria sanguinalis, Aeschynomene indica, and Abutilon avicennae*, and > 90% control of *Echinochlia crus-galli, Panicum dichotomiflorum, Xanthium strumarium, and Calystegia japonica*. Herbicidal activity against *Agropyron smithii* was slightly weaker (Table 1).

**Table 1.** Herbicidal activity of foliar application of methanol extracts from *Ailanthus altissima* to several weeds in a greenhouse condition.

| Rates<br>(mg/mL) | Herbicidal activity (%) |  |  |  |  |  |  |  |  |  |
| --- | --- | --- | --- | --- | --- | --- | --- | --- | --- | --- |
|  | SOR | ECHC | AGRS | DIGS | PAN | SOL | AESI | ABUT | XAN | CAG |
| 10 | 100 | 95 | 30 | 100 | 98 | 70 | 100 | 100 | 90 | 95 |
| 5 | 40 | 40 | 20 | 100 | 70 | 60 | 100 | 80 | 40 | 70 |
| 2.5 | 40 | 30 | 10 | 90 | 20 | 20 | 98 | 98 | 40 | 70 |
| 1 | 40 | 30 | 10 | 70 | 20 | 20 | 95 | 98 | 40 | 70 |

SORBI ; *Sorghum bicolor*, ECHCG ; *Echinochloa crus-galli*, AGRSM ; *Agropyron smithii*, DIGSA ; *Digitaria sanguinalis*, PANDI ; *Panicum dichotomiflorum*, SOLNI ; *Solanum nigrum*, AESIN ; *Aeschynomene indica*, ABUTH ; *Abutilon avicennae*, XANSI ; *Xanthium strumarium*, CAGHE ; *Calystegia japonica*.

The main characteristic was its quick action upon foliar application. External symptoms began to appear within 24 h of exposure, and the weeds were completely controlled five days after treatment. MeOH extract showing herbicidal activity was purified by ethyl acetate fractionation and a series of chromatographic techniques, including silica gel column, C18 Sep-pak cartridge, Sephadex LH20, and preparative high-performance liquid chromatography (HPLC). Four quassinoids, **1-4** (Figure 1, Figs. S1∼4), were isolated from the MeOH extract of *Ailanthus altissima* leaves and showed herbicidal activity of 98, 95, 85, and 75%, respectively, when applied to *D. sanguiinalis* at a concentration of 10 μg/mL (Table 2, Figure 2).

**Table 2.** Herbicidal activity of isolated compounds against *Digitaria sanguinalis*.

|  | Herbicidal activity at 10 µg/mL (%) |
| --- | --- |
| Compound 1 | 98 |
| Compound 2 | 95 |
| Compound 3 | 85 |
| Compound 4 | 75 |

**Fig 1.**
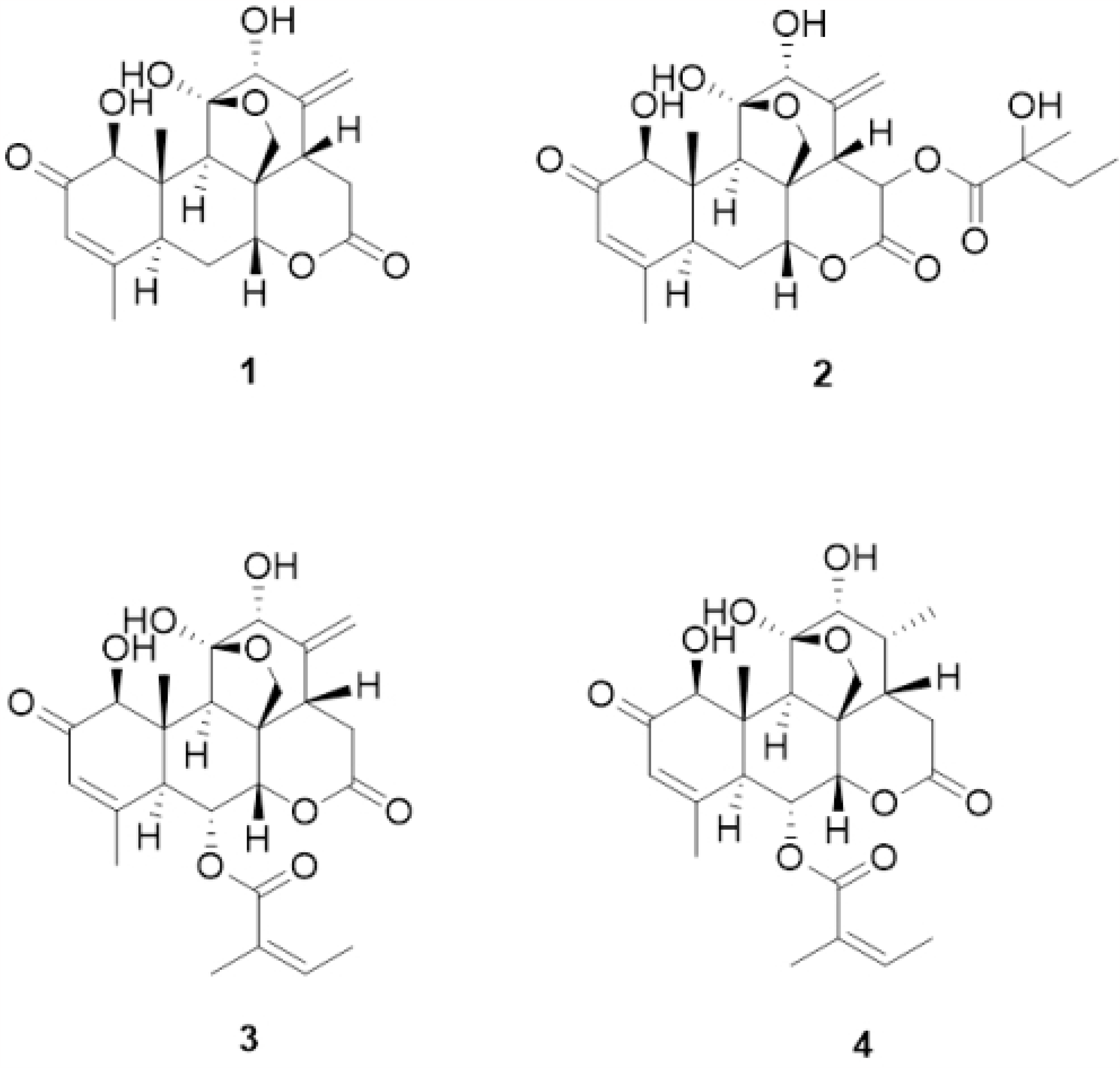
Structures of compounds 1-4.

**Fig 2.**
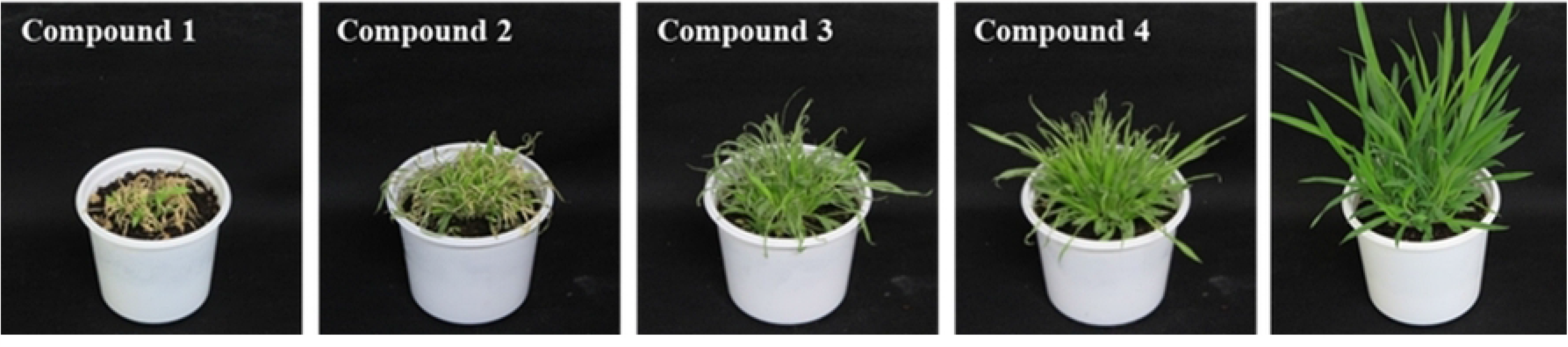
Herbicidal activity against *Digitaria sanguinalis* at same concentration of 10 μg/mL.

Among these, **3** was a new compound, and the characterization and spectroscopic analysis of this compound were performed by data comparison with literature values[14]. The ^1^H and ^13^C NMR (nuclear magnetic resonance) data for these compounds are shown in the experimental section. The ^1^H (Table 3) and ^13^C NMR (Table 4) signals assignments were aided by ^1^H-^1^H COSY (correlation spectroscop Y), HSQC (heteronuclear single quantum correlation) and HMBC (heteronuclear multiple bond correlation) experiments. The other compounds were identified as ailanthone (**1**)[15], 13, 18-dehydroglaucarubinone (**2**)[15,16], and 6-α-tigloyloxychaparrinone **(4)[**17,18].

**Table 3.** ^1^H-NMR Data (DMSO-*d*_6_) for compounds 1-4.

|  | Compound |  |  |  |
| --- | --- | --- | --- | --- |
| Proton | 1 | 2 | 3 | 4 |
| H-1 | 4.28(s) | 4.42 (d, 2.1) | 4.39 (s) | 4.30 (d, 2.8) |
| H-3 | 5.99 (d, 0.9) | 5.99 (q, 0.7) | 6.02 (s) | 6.0 (s) |
| H-5 | 2.88 (d, 10.8) | 3.08 (d, 11.4) | 3.46 (d, 11.9) | 3.39 (d, 11.9) |
| H-6 | 2.03 (m) | 2.06 (m) | 5.49 (dd, 11.9, 2.8) | 5.49 (dd, 11.9, 2.8) |
| H-7 | 4.55 (t, 2.8) | 4.70 (t, 2.8) | 4.63 (d, 2.8) | 4.56 (d, 2.8) |
| H-9 | 2.81 (s) | 2.97(s) | 2.83(s) | 2.61(s) |
| H-12 | 3.67 (d, 3.6) | 3.67 (d, 4.9) | 3.69 (d, 3.5) | 3.21 (t, 4.9) |
| H-13 | - | - | - | 20.9 (m) |
| H-14 | 2.77 (dd, 13.5, 5.4) | 3.04 (d, 11.4) | 2.85 (dd, 13.3, 5.6) | 2.05 (m) |
| Ha-15 | 2.98 (dd, 18.4, 13.5) | 5.64 (br s) | 3.01 (dd, 18.4, 13.3) | 2.72 (dd, 18.2, 13.3) |
| Hb-15 | 2.44 (dd, 18.4, 5.4) | - | 2.47 (dd, 18.4, 5.6) | 2.40 (dd, 18.2, 5.6) |
| H-18 | 1.93 (s) | 1.93 (s) | 1.94 (s) | 1.94 (s) |
| H-19 | 1.06 (s) | 1.06 (s) | 1.21 (s) | 1.22 (s) |
| Ha-20 | 3.81 (d, 9.0) | 3.81 (d, 8.4) | 3.91 (d, 8.4) | 3.93 (d, 8.4) |
| Hb-20 | 3.27 (d, 9.0) | 3.32 (d, 8.4) | 3.39 (d, 8.4) | 3.64 (d, 8.4) |
| Ha-21 | 5.05 (d, 0.9) | 5.09 (d, 1.4) | 5.02 (d, 1.4) | - |
| Hb-21 | 5.02 (d, 0.9) | 4.99 (br s) | 5.01 (br s) | - |
| Ha-3' | - | 1.70 (dq, 14.0, 7.0) | 6.91 (dq, 7.7, 2.1) | 6.91 (dq, 7.7, 2.1) |
| Hb-3' | - | 1.53 (dq, 14.0, 7.0) | - | - |
| H-4' | - | 0.81 (t, 7.0) | 1.823 (d, 4.2) | 1.81 (d, 7.7) |
| H-5' | - | 1.30 (s) | 1.829 (s) | 1.82 (br s) |
| 1-OH | 7.08 (s) | 7.16 (d, 2.1) | 7.28 (s) | 7.19 (d, 2.8) |
| 11-OH | 8.23 (s) | 8.33 (s) | 8.15 (s) | 8.0 (s) |
| 12-OH | 5.40 (d, 4.5) | 5.35 (br s) | 5.55 (d, 4.2) | 5.14 (d, 4.9) |

**Table 4.** ^13^C-NMR Data (DMSO-*d*_6_) for compounds 1-4.

| Carbon | Compound |  |  |  |
| --- | --- | --- | --- | --- |
|  | 1 | 2 | 3 | 4 |
| C-1 | 82.4 | 82.1 | 82.4 | 82.5 |
| C-2 | 197.1 | 197.0 | 196.6 | 196.5 |
| C-3 | 125.0 | 124.8 | 127.6 | 127.5 |
| C-4 | 162.4 | 162.4 | 161.1 | 161.2 |
| C-5 | 41.2 | 40.7 | 44.1 | 44.1 |
| C-6 | 25.1 | 24.6 | 67.3 | 67.3 |
| C-7 | 77.5 | 77.8 | 77.6 | 77.6 |
| C-8 | 44.5 | 46.4 | 45.2 | 45.7 |
| C-9 | 43.3 | 44.0 | 42.0 | 41.8 |
| C-10 | 44.5 | 44.5 | 47.2 | 47.1 |
| C-11 | 108.8 | 108.6 | 109.0 | 109.1 |
| C-12 | 79.0 | 78.7 | 79.1 | 78.1 |
| C-13 | 146.6 | 142.4 | 146.1 | 30.2 |
| C-14 | 46.1 | 50.2 | 46.1 | 40.6 |
| C-15 | 34.3 | 68.2 | 34.0 | 29.3 |
| C-16 | 169.1 | 166.7 | 168.2 | 168.9 |
| C-17 | 22.4 | 22.2 | 24.7 | 24.5 |
| C-18 | 9.5 | 9.5 | 10.9 | 10.8 |
| C-19 | 71.1 | 70.7 | 70.2 | 69.3 |
| C-20 | 117.6 | 119.8 | 117.5 | 12.5 |
| C-1' | - | 174.2 | 165.9 | 165.8 |
| C-2' | - | 73.9 | 127.9 | 127.8 |
| C-3' | - | 32.8 | 139.2 | 139.0 |
| C-4' | - | 8.0 | 14.4 | 14.3 |
| C-5' | - | 25.8 | 11.9 | 11.8 |

Compound **3** was obtained as a white solid. The molecular formula was established as C_25_H_32_O_9_ based on a quasi-molecular ion at m/z 497.1785 [M + Na]^+^ (calcd 497.1788) in its HRTOFMS. The ^1^H NMR spectrum of **3** displayed signals for two olefinic protons [δ_H_ 6.91 (1H, dq, *J* = 7.7, 2.1, H-3’), 6.02 (1H, s, H-3)], an exo-methylene [δ_H_ 5.02 (1H, d, *J* = 2.1, H_a_-21), 5.01 (1H, s, H_b_-21)], four oxygenated methines [δ_H_ 5.49 (1H, dd, *J* = 11.9, 2.8, H-6), 4.63 (1H, d, *J* = 2.8, H-7), 4.39 (1H, s, H-1), 3.69 (1H, d, *J* = 3.5, H-12)], one oxygenated methylene [δ_H_ 3.91 and 3.39 (each 1H, d, *J* = 8.4, H-20)], one methylene proton [δ_H_ 3.01 (1H, dd, *J* = 18.4, 13.3, H_a_-15) and 2.40 (1H, dd, *J* = 18.4, 5.6, H_b_-15)] and four methyl groups [δ_H_ 1.94 (3H, s, H-18), 1.829 (3H, br s, H-5’), 1.823 (3H, d, *J* = 7.7, H-4’), and 1.21 (3H, s, H-19)] (Fig S9). The ^13^C NMR spectrum exhibited 25 carbon signals, including three carbonyl signals, six olefinic carbon signals, one hemiketal carbon signal, four methyl carbon signals, two methylene carbon signals, seven methine carbon signals, and two quaternary carbon signals(Fig S10). Comparison of the NMR data of **3** with those of ailanthone (**1**) revealed that the two compounds possessed the same quassinoid moiety, except that the C6 position in ailanthone (**1**) was substituted with a tigloyloxy group in **3**. The ^1^H-^1^H COSY between H-3’ (δ_H_ 6.91) and H-4’ (δ_H_ 1.823) and the HMBC correlation between H-3’ (δ_H_ 6.91), H-5’ (δ_H_ 1.829), and C1’ (δ_C_ 165.8) and H-4’ (δ_H_ 1.81) and C-2’ (δ_C_ 127.8) confirmed the existence of a tigloyloxy group and that it was connected to the C6 position of **3**, established by a long-range correlation between δ_H_ 5.49 (H-6 of quassinoid moiety) and carboxylic carbon (δ_C_ 165.9, C1’ of tigloyloxy group) in the HMBC spectrum of **3**(Figs. S11, 12, 13). The correlations between H-9 and H-1/H-5 and between H-7 and H-14 in the ROESY spectrum(Fig S14) suggested that the configuration of **3** was the same as that of ailanthone (**1**). And the ROESY correlation between H-3’ and H-5’ indicated that the geometry of the double-bond of the tigloyloxy group was Z-form. Thus, the structure of compound **3** was determined to be 6-α-tigloyloxyailanthone. The relative stereochemistry of **3** was confirmed from its 2D-ROESY NMR spectrum. The NOE correlations are shown by arrows in Figure 3 (2D-ROESY).

**Fig 3.**
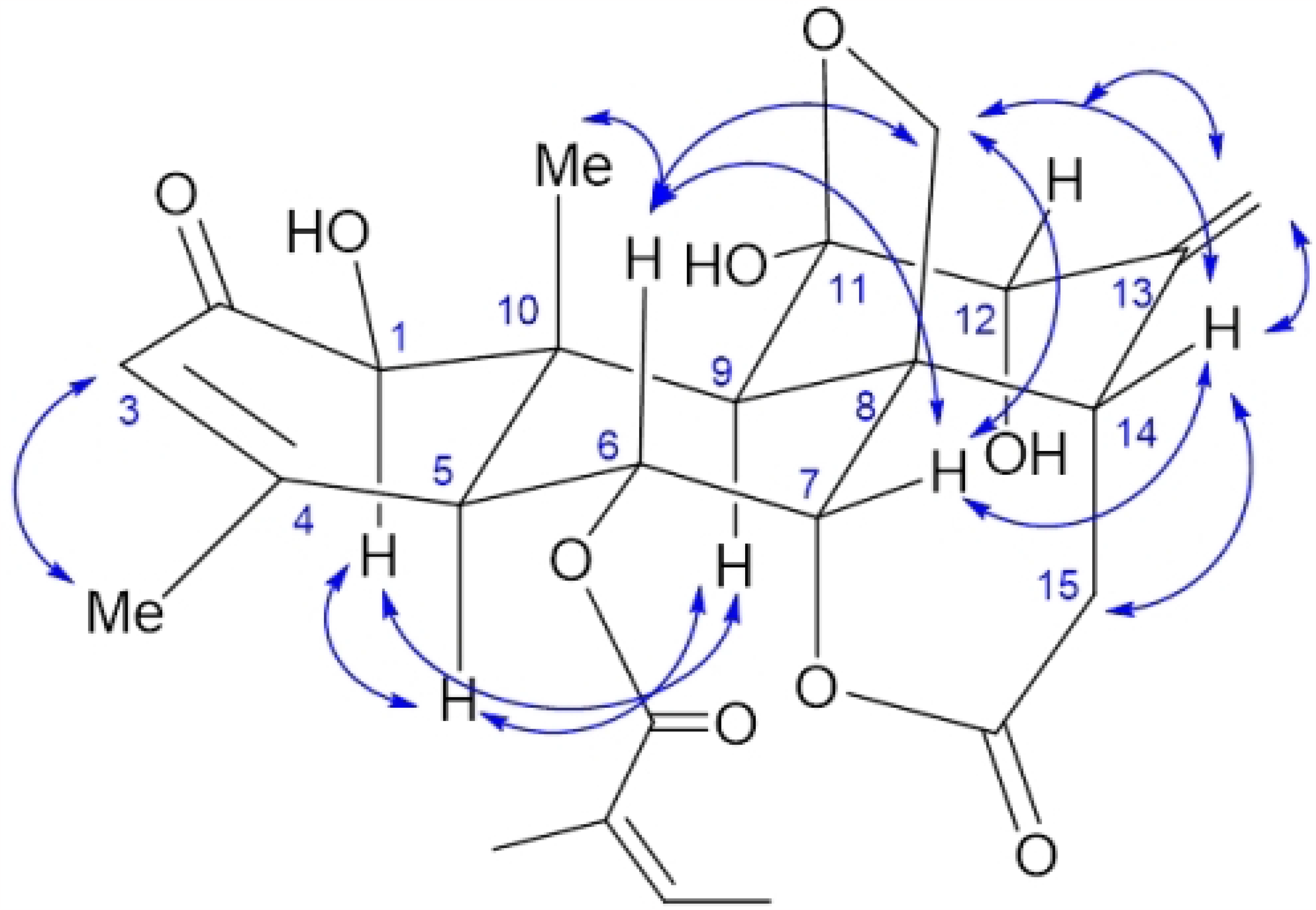
2D ROESY (NOE) correlation of 3.

## Acknowledgment

This work was carried out with the support of the Cooperative Research Program for Agriculture Science & Technology Development (Project No. PJ0161282023), Rural Development Administration, Republic of Korea.

## Supporting Information

Structures and spectra of compound 1, 2, 3 and 4.

## References

1. Polonsky J. Quassinoid Bitter Principles. In: Grisebach H, Kirby GW, Herz W, editors. Fortschritte Der Chemie Organischer Naturstoffe: Progress in the Chemistry of Organic Natural Products. Vienna: Springer Vienna; 1973, pp 101–102.

2. Curcino Vieira IJ, Braz-Filho R. Quassinoids: Structural Diversity, biological activity and Synthetic Studies. Stud Nat Prod Chem. 2006; 433–492. doi:10.1016/s1572-5995(06)80032-3

3. Guo Z, Vangapandu S, Sindelar R, Walker L, Sindelar R. Biologically active quassinoids and their chemistry: Potential leads for Drug Design. Curr Med Chem. 2005;12: 173–190. doi:10.2174/0929867053363351

4. Alves IABS, Miranda HM, Soares LAL, Randau KP. Simaroubaceae family: Botany, chemical composition and Biological Activities. Rev Bras Farmacogn. 2014;24: 481–501. doi:10.1016/j.bjp.2014.07.021

5. Polonsky J. Quassinoid Bitter Principles II. In: Herz W, Grisebach H, Kirby GW, Tamm C, editors. Fortschritte der Chemie organischer Naturstoffe / Progress in the Chemistry of Organic Natural Products. Vienna: Springer Vienna; 1985, pp 221–264.

6. Fukamiya N, Okano M, Miyamoto M, Tagahara K, Lee K-H. Antitumor agents, 127. Bruceoside C, a new cytotoxic quassinoid glucoside, and related compounds from Brucea javanica. J Nat Prod. 1992;55: 468–475. doi:10.1021/np50082a011

7. Ang H, Chan K, Mak J. In vitro antimalarial activity of quassinoids from Eurycoma longifoliaagainst Malaysian chloroquine-resistant Plasmodium falciparum Isolates. Planta Med. 1995;61: 177–178. doi:10.1055/s-2006-958042

8. Apers S, Cimanga K, Vanden Berghe D, Van Meenen E, Longanga AO, Foriers A, et al. Antiviral activity of Simalikalactone D, a Quassinoid from Quassia Africana. Planta Med. 2002;68: 20–24. doi:10.1055/s-2002-19870

9. Silva RL, Lopes AH, França RO, Vieira SM, Silva EC, Amorim RC, et al. The quassinoid isobrucein B reduces inflammatory hyperalgesia and cytokine production by post-transcriptional modulation. J Nat Prod. 2015;78: 241–249. doi:10.1021/np500796f

10. Leskinen V, Polonsky J, Bhatnagar S. Antifeedant activity of quassinoids. J Chem Ecol. 1984; 10: 1497–1507.

11. Daido M, Fukamiya N, Okano M, Taoahara K, Hatakoshi M, Yamazaki H. Antifeedant and insecticidal activity of quassinoids against diamondback moth (plutella xylostella). Biosci Biotechnol Biochem. 1993;57: 244–246. doi:10.1271/bbb.57.244

12. Tada H, Yasuda F, Otani k, Doteuchi M, Ishihara Y, Shiro M. Cheminform abstract: New Antiulcer quassinoids from Eurycoma longifolia. ChemInform. 1991;22. doi:10.1002/chin.199132267

13. Heisey R, Kish Heisey T. Herbicidal effects under field conditions of Ailanthus altissima bark extract, which contains ailanthone. Plant Soil. 2003;256: 85–99. doi:10.1023/a:1026209614161

14. Kubota K, Fukamiya N, Tokuda H, Nishino H, Tagahara K, Lee K-H, et al. Quassinoids as inhibitors of Epstein-Barr virus early antigen activation. Cancer Lett. 1997;113: 165–168. doi:10.1016/s0304-3835(97)04607-7

15. Kubota K, Fukamiya N, Hamada T, Okano M, Tagahara K, Lee K-H. Two new quassinoids, Ailantinols A and B, and related compounds from ailanthus altissima. J Nat Prod. 1996;59: 683–686. doi:10.1021/np960427c

16. Polonsky J, Varon Z, Jacquemin H, Pettit GR. The isolation and structure of 13,18-dehydroglaucarubinone, a new antineoplastic quassinoid fromsimarouba Amara. Experientia. 1978;34: 1122–1123. doi:10.1007/bf01922904

17. Carter CA, Tinto WF, Reynolds WF, McLean S. Quassinoids from quassia multiflora: Structural assignments by 2D NMR spectroscopy. J Nat Prod. 1993;56: 130–133. doi:10.1021/np50091a019

18. Okunade AL, Bikoff RE, Casper SJ, Oksman A, Goldberg DE, Lewis WH. Antiplasmodial activity of extracts and quassinoids isolated from seedlings ofAilanthus altissima (Simaroubaceae). Phytother Res. 2003;17: 675–677. doi:10.1002/ptr.1336

